# Effects of a native, dominant tree, *Colophospermum mopane*, on diversity of plants, insects, and vertebrates in South African savannas

**DOI:** 10.1101/2025.11.17.688835

**Authors:** Fernando P. Gaona, Tomáš Albrecht, Jan Čuda, Sylvain Delabye, Llewellyn C. Foxcroft, Valeriy Govorov, Martin Hejda, Ivan Horáček, Sandra MacFadyen, Pavel Potocký, Klára Pyšková, Vladimír Remeš, Ondřej Sedláček, Markéta Staňková, David Storch, Petr Pyšek, Robert Tropek

**Author notes:** Corresponding authors (FPG); (RT).

## Abstract

Afrotropical savannas are biodiversity-rich ecosystems increasingly threatened by woody plant encroachment. In southern Africa, the leguminous tree *Colophospermum mopane* dominates over one-third of the savanna region and is projected to expand substantially under climate change. Yet, the consequences of its local dominance for biodiversity remain poorly understood. We conducted the first landscape-scale, multi-taxon assessment of mopane’s bottom-up effects, analysing species richness and community composition of vascular plants, insects, birds, bats, and non-flying mammals across a gradient of mopane cover in Kruger National Park, South Africa. Our replicated plot-based study found that species richness of birds, mammals, bats, and insects declined significantly with increasing mopane dominance, with the steepest reductions in birds. Plant overall species richness was unaffected, although grasses showed a weak positive trend. We also revealed significant community compositional shifts in birds, bats, and mammals, while insect communities lost species without systematic composition change. Functional-group analyses confirmed species richness declines for herbivores across taxa, and for bird carnivores and omnivores, pointing to mopane’s role as a strong ecological filter that reduces host plant and other resources availability, with consequences to the higher trophic levels. These results highlight mopane dominance as a potential driver of biodiversity simplification in African savannas, with cascading implications for ecosystem functioning. Given projections of mopane expansion and its socioeconomic value to local communities, management and policy must avoid promoting mopane dominance in land-use or restoration schemes. Safeguarding heterogeneous savanna mosaics will be essential for conserving biodiversity and ecosystem resilience under climate change.

## Introduction

Afrotropical savannas rank among the most species-rich terrestrial biomes, harbouring a plethora of species adapted to this highly dynamic ecosystem shaped by interacting gradients of climate, soil, fire, and herbivory (Parr et al., 2014; Nerlekar & Veldman, 2020). Despite the high environmental heterogeneity in this part of Africa resulting in a wide variety of savanna types, a leguminous tree species, *Colophospermum mopane* (commonly known as mopane), occurs across more than one-third (> 500,000 km^2^) of the southern African savanna region (Mapaure, 1994; Makhado et al., 2014; Stevens, 2021). Within this area, mopane often dominates local vegetation and, under suitable conditions, can form vast, contiguous woodlands (Timberlake, 1995; Makhado et al., 2014). Moreover, recent modelling studies have projected a 20–100 % expansion of its range under future climate scenarios, identifying mopane as a likely “regional winner” in warming savannas (Kapuka et al., 2022; Jinga et al., 2023). Such dominance and prospective spread prompt questions about its effects on the biodiversity of Afrotropical savannas.

The dominance of mopane in some of the hottest, driest, and most nutrient-poor parts of the Afrotropical savanna is enabled by a suite of complementary adaptations (Makhado et al., 2014; Stevens, 2021). Its high heat tolerance is mainly linked to reflective cuticles and efficient stomatal control (Timberlake, 1995; Stevens et al., 2014). Dense fine-root networks confined to upper soil layers enable rapid uptake of water and nutrients after sporadic rains (Smit & Rethman, 1998). Although lacking typical nodules, mopane associates with nitrogen-fixing bacteria in its rhizosphere, probably gaining supplemental N on infertile soils (Burbano et al., 2015). After damage, vigorous basal resprouting quickly restores the canopy, and calcium-oxalate–rich wood increases fire resistance (Mlambo & Mapaure, 2006; Makhado et al., 2014). Its drought-deciduous leaves flush after the first rains and are shed in the dry season, while osmotic regulation helps maintain tissue hydration during dry intra-seasonal periods (Timberlake, 1995; Makhado et al., 2018). Mopane foliage combines high protein content with elevated tannin and phenolic levels, deterring herbivores, and its coppicing growth after damage produce multi-stem clusters of hard branches keeping its shoots out of reach by many medium and large herbivores (Wessels et al., 2007; Smallie & O’Connor, 2000). Together, these traits provide a strong advantage under fire, heat, drought, and nutrient stress.

Some of mopane’s adaptations may also reshape local environmental conditions. Its tannin- and lignin-rich leaves decompose slowly, accumulating in a thick litter layer that elevates soil pH and electrical conductivity, reduces evaporation, and physically suppresses seedling emergence (Mlambo & Mwenje, 2010; Smit & Rethman, 1998). At the same time, mopane contributes to nutrient cycling by annually returning calcium, potassium, and nitrogen via litterfall (Mlambo & Nyathi, 2008), and it may enrich surface soils through symbioses with nitrogen-fixing bacteria (Burbano et al., 2015). Its shallow root system rapidly extracts moisture from the upper soil layers following rainfall (Smit & Rethman, 1998; Makhado et al., 2014). Physically, vigorous post-fire and post-browsing resprouting produces a low, multi-stem coppice, especially in areas of high elephant activity, reducing vegetation complexity and altering fire regimes (Mlambo & Mapaure, 2006; Smallie & O’Connor, 2000). Under suitable conditions, these processes generate structurally and microclimatically distinct patches of shaded, litter-rich mopane woodland that contrast sharply with the surrounding mixed savanna (Timberlake, 1995). Such changes are likely to influence other organisms occupying these habitats.

Direct and indirect influences of mopane on syntopic species have been described, although comprehensive data and detailed studies remain scarce. Due to its effects on environmental conditions, shade-intolerant herbs and tree seedlings are expected to decline beneath its canopy (Timberlake, 1995; Mlambo et al., 2005; Stevens, 2021). Allelopathy by mopane has been repeatedly proposed (e.g. Stevens, 2021; Georginah & Maanda, 2015) but remains untested. Insects show a contrasting pattern: a few specialised herbivores, such as the saturniid “mopane worm” (*Gonimbrasia belina*) and various sap-sucking hemipterans, thrive on the tree’s protein-rich yet chemically defended foliage, while generalist herbivore richness appears exceptionally limited (Timberlake, 1995), although detailed studies are lacking. Large vertebrate browsers often exploit seasonal windows of reduced defence, with elephants, kudu, and impala relying on mopane leaves and twigs during the dry season, when tannin concentrations are lower (Ben-Shahar, 1996; Styles & Skinner, 2000; Makhado et al., 2018). Among birds, some cavity-nesting species, such as Lilian’s lovebird, depend on hollows in mature mopane trees (Mzumara et al., 2019), and some insectivorous birds feed on its insect herbivores (Herremans et al., 1995). Similar effects can be expected for bats, but no studies have yet addressed this. Thus, mopane provides key resources for a limited community of adapted specialists while imposing strong resource and habitat filters on many other co-occurring species.

Unfortunately, such effects of mopane dominance on species communities remain poorly understood. To date, only two small-scale studies have addressed plant communities, both conducted in the Makuya Nature Reserve, South Africa, and with conflicting results. Khavhagali and Ligavha-Mbelengwa (2009) found reduced species richness under mopane canopies on loamy soils but the opposite trend on sandy soils, while Georginah and Maanda (2015) reported lower biomass and species richness in mopane stands compared to open savannas. Both surveys were spatially limited, lacked replication across soil types, and, crucially, contrasted mopane woodland with open savanna rather than with mixed woodland, thereby conflating mopane dominance with mixed tree cover. Additionally, an experimental study showed an increased grass biomass after repeated small-scale mopane removal, in comparison with control mopane-dominated plots (Wedel et al., 2024). To our knowledge, no studies have examined how insect, bird, bat, or mammal communities vary along a gradient of mopane dominance. Vertebrate data remain restricted to individual species’ habitat use (e.g. Ben-Shahar, 1996; Wedel et al., 2024) or non-comparative faunal surveys over broader areas within the mopane’s distribution range. Consequently, we lack any systematic, multi-taxon, landscape-scale assessment of how mopane cover influences species richness and community composition. Without such knowledge, we cannot adequately evaluate mopane’s role in shaping savanna biodiversity, nor can we anticipate the ecological consequences of its projected expansion under climate change.

Here we present the first landscape-scale, multi-taxon study examining how mopane dominance affects savanna biodiversity. It integrates data on vascular plants, insects, birds, bats, and non-flying mammals across a gradient of mopane cover in Kruger National Park, a key biodiversity hotspot within the Afrotropical savanna. The study aims to quantify how increasing mopane cover influences species richness and community composition across major taxonomic and trophic groups. We predict that species richness will generally decline with increasing mopane cover, and that community composition will shift significantly along the cover gradient. Nevertheless, we expect the magnitude and direction of these responses will differ among groups. Species richness of woody plants and herbs is predicted to decline due to competitive exclusion and habitat alteration, while grasses may remain stable or benefit from reduced competition and nutrient enrichment under the mopane canopy. Insect herbivores are expected to decline with decreasing host-plant diversity, with cascading negative effects on insectivorous insects, birds, and bats. Omnivores may also respond negatively to reduced habitat complexity and lower food diversity. In contrast, mammalian herbivores may show neutral or even positive responses due to mopane’s dry-season forage value, and their predators are therefore unlikely to be significantly affected. By filling the crucial data gaps, this study provides essential insights into mopane’s role in structuring communities and informs conservation strategies in light of its projected range expansion.

## Methods

### Study species

*Colophospermum mopane* (Kirk ex Benth.) J. Léonard is a leguminous tree or multi-stemmed shrub with distinctive bifoliolate, butterfly-shaped leaves that dominates roughly one-third of the southern-African savanna zone between 9° S and 25° S (Mapaure, 1994; Makhado et al., 2014; Fig. 1a). Growth form varies with site conditions: stunted “mopane scrub” ≤ 2–4 m tall occurs on very dry, nutrient-poor soils, whereas well-watered alluvial flats support single-stem trees up to 20–24 m in height (Timberlake, 1995). The species thrives in areas receiving ∼400– 800 mm annual rainfall and mean summer temperatures > 30 °C (Stevens, 2021). Owing to its dominance and drought-hardiness, mopane is a key resource in rural economies, valued for durable timber, firewood, and livestock forage, as well as supporting the seasonal harvest of edible saturniid caterpillars known as mopane worms (Timberlake, 1995; Makhado et al., 2012).

**Figure 1.**
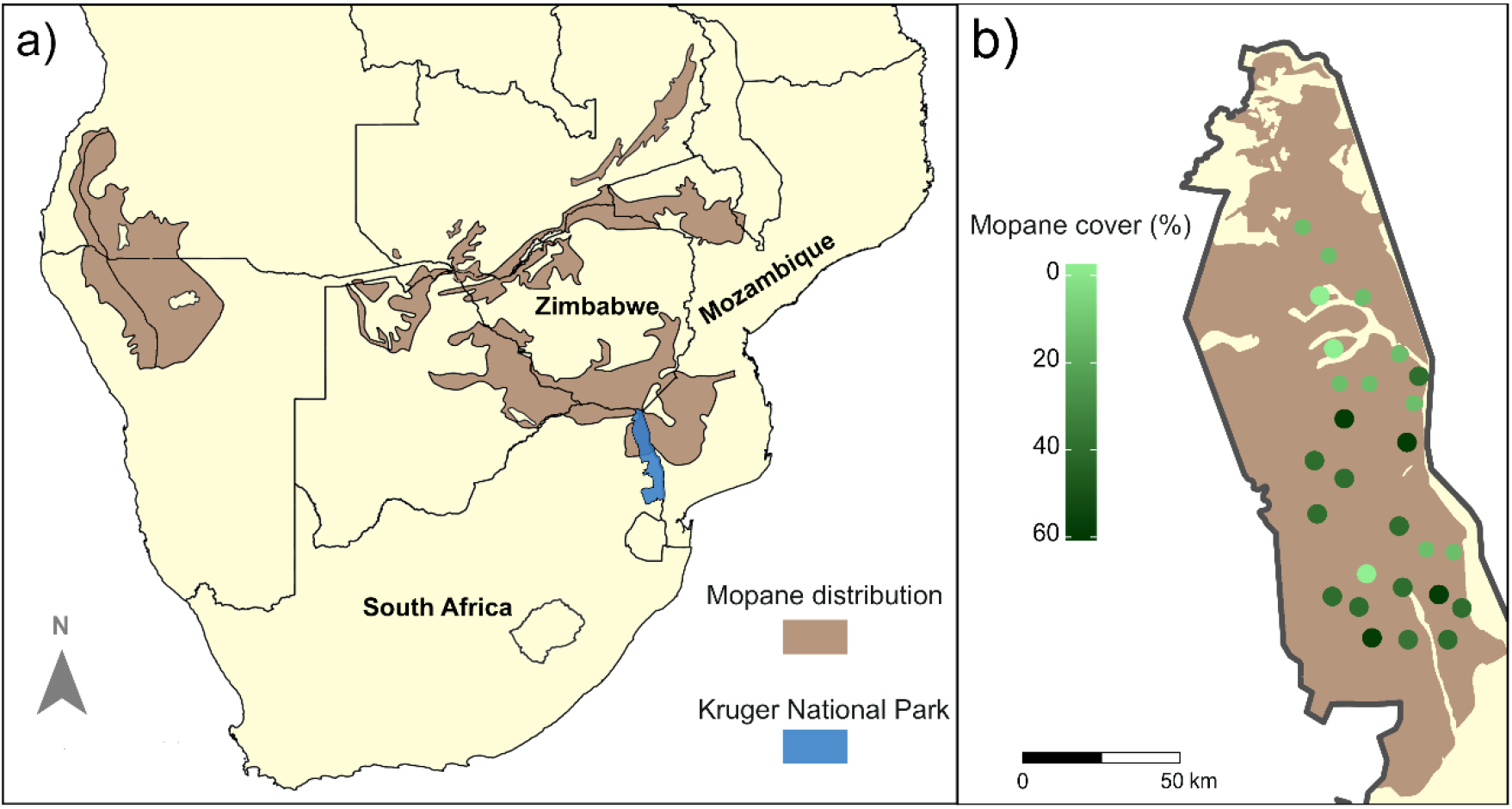
a) Distribution of mopane (*Colophospermum mopane*) and Kruger National Park (KNP) in southern Africa (modified after Stevens, 2021). b) Distribution of the 27 sampling plots within the mopane range in KNP. Circle colors indicate the percentage of mopane cover at each plot.

### Study area and sampling design

The study was carried out in the northern part of Kruger National Park (KNP), South Africa (Fig. 1b). Established in 1926, KNP is one of the largest and oldest protected areas in the region, covering approximately 20,000 km^2^. This region falls predominantly within a semi-arid tropical and subtropical climatic zones and receives relatively low annual precipitation (∼450 mm), compared to the southern parts of KNP (Chadwick et al., 2013). Elevation ranges from 140 to 400 m a.s.l., and the landscape is intersected by major rivers, including Letaba, Luvuvhu, and Shingwedzi (du Toit et al., 2003). Large parts of northern KNP are dominated by mopane woodlands (*Colophospermum mopane*), one of the most abundant of the park’s > 400 woody species (Van Wyk, 2023).

We used biodiversity datasets sampled within the MOSAIK project (Monitoring Savanna Biodiversity in the Kruger National Park), which aims to disentangle effects of key drivers shaping spatiotemporal dynamics of savanna biodiversity across trophic levels (e.g. Pyšek et al., 2020; Hejda et al., 2022; Čuda et al., 2024; Gaona et al., 2025). The data were collected at 27 plots (50 × 50 m) distributed across the mopane-distribution area in KNP (Fig. 1b). The plots were organised into nine triplets originally established along a gradient of water availability (near perennial rivers, near seasonal rivers, and far from water sources), covering local environmental heterogeneity (Hejda et al., 2022; Gaona et al., 2025). Within each triplet, plots were selected within the same landsystem and bedrock type, and with similar vegetation structure and density. Triplets also structured the sampling design of some groups: moths and bats were sampled on the same night and birds in the same morning across the three plots of a triplet. Across the 27 plots, mopane cover ranged from 0% to 62.5% (mean ±SD: 29.54 ±19.17%; Fig. 1b).

### Diversity data and functional groups

We analysed species richness and community composition of five taxonomic groups: vascular plants (hereinafter referred to as plants), insects (moths and mantises), bats, medium- and large-sized non-flying mammals (hereinafter referred to as mammals), and birds; we used datasets from previous studies for all groups (Hejda et al., 2022; Gaona et al., 2025; Staňková et al., 2024; Čuda et al., 2024), except for birds. Plants were surveyed during the wet season (details in Hejda et al., 2022); insects were sampled using portable automated light traps during the early and peak wet seasons (details in Gaona et al., 2025); bats were recorded with automated ultrasound detectors on the same nights as insect sampling (details in Staňková et al., 2024); and mammals were monitored with camera traps during both dry and wet seasons (details in Čuda et al., 2024). Birds were surveyed within a 100-m radius from each plot centre during morning hours using the point-count method (Bibby et al., 2000). Each point was surveyed twice during the wet season (January and March), with birds recorded during four consecutive 10-min intervals per visit (40 min per visit, 80 min in total). For analyses, we used the maximum abundance of each species detected across any of the 10-min intervals.

All recorded species were assigned to functional groups based on growth form (plants: grasses, herbs, and woody species) or trophic level (animals: herbivores, carnivores, omnivores). All moths were classified as herbivores, and all mantises as carnivores. Birds were assigned to trophic groups according to the *Trophic*.*Level* trait in the *AVONET* database (Tobias et al., 2022), and mammals according to their prevailing diet in the *MammalDIET* database (Kissling et al., 2014). Bats were not included in functional group analyses, as only insectivorous species were recorded (Staňková et al., 2024). Each species was assigned to a single functional group.

### Statistical analyses

All analyses were conducted in R v. 4.3.3 (R Core Team 2024). To evaluate the effects of mopane cover (logit-transformed) on species richness across taxonomic and functional groups, we fitted Generalized Linear Mixed Models (GLMMs), with triplet included as a random intercept to account for spatial clustering of plots and the sampling design of some groups. Separate models were built for each taxonomic group (plants, insects, bats, mammals, birds) and for the functional groups nested within them (plants: grasses, herbs, and woody species; each animal group: herbivores, carnivores, and omnivores). An initial visual inspection of the data assumed a Poisson error distribution. Dispersion was assessed using simulation-based diagnostics and DHARMa dispersion tests (Hartig, 2022). Overdispersed models were re-fitted with a negative binomial distribution, while underdispersed models were estimated using quasi-Poisson error structure (Table 1). All GLMMs were implemented with the *lme4* (Bates et al., 2015) and *glmmTMB* (McGillycuddy et al., 2025) packages. Specifically, Poisson and quasi-Poisson models were fitted with *lme4*, whereas negative binomial models were fitted using *glmmTMB*. To facilitate comparison of effect magnitudes across groups, standardized regression coefficients were calculated using the *effectsize* package (Ben-Shachar et al., 2021).

**Table 1.** Results of generalized linear mixed models (GLMMs) testing the effect of mopane cover (logit-transformed) on species richness of taxonomic groups and their respective functional groups in Kruger National Park, South Africa. For each model, the error distribution, estimated regression coefficient (Estimate), standard error (Std. Error), and p-value are reported. Statistically significant effects (p < 0.05) are highlighted in bold. See Fig. 2 for results visualisation.

| **Taxonomic group** | **Functional group** | **Model distribution** | **Estimate** | **Std.Error** | **p-value** |
| --- | --- | --- | --- | --- | --- |
| Bats | *all species* | Quasi-Poisson | -0.021 | 0.010 | **0.041** |
| Birds | *all species* | Negative Binomial | -0.103 | 0.046 | **0.013** |
|  | *herbivores* | Poisson | -0.094 | 0.029 | **0.001** |
|  | *carnivores* | Negative Binomial | -0.102 | 0.039 | **0.008** |
|  | *omnivores* | Poisson | -0.111 | 0.054 | **0.042** |
| Mammals | *all species* | Poisson | -0.066 | 0.022 | **0.003** |
|  | *herbivores* | Poisson | -0.072 | 0.029 | **0.011** |
|  | *carnivores* | Poisson | -0.029 | 0.091 | 0.754 |
|  | *omnivores* | Poisson | -0.053 | 0.040 | 0.183 |
| Insects | *all species* | Poisson | -0.040 | 0.103 | **<0.001** |
|  | *herbivores* | Poisson | -0.042 | 0.010 | **<0.001** |
|  | *carnivores* | Poisson | 0.029 | 0.043 | 0.500 |
| Plants | *all species* | Poisson | -0.000 | 0.014 | 0.992 |
|  | *grasses* | Poisson | 0.053 | 0.033 | 0.071 |
|  | *herbs* | Poisson | -0.001 | 0.020 | 0.935 |
|  | *woody* | Poisson | -0.027 | 0.026 | 0.291 |

To assess changes in community composition along the gradient of mopane cover, we applied distance-based redundancy analysis (db-RDA) using Bray–Curtis dissimilarity matrices in the *vegan* package (Oksanen et al., 2022). For each taxonomic group, the dissimilarity matrix based on species abundances at individual plots was regressed on logit-transformed mopane cover. We used a split-plot permutation design (9999 permutations), stratifying permutations to constrain swaps within triplets, thereby accounting for the hierarchical sampling structure while testing the effects of mopane cover.

## Results

Across the 27 sampling plots, the compiled datasets included 357 plant species (185 herbs, 76 grasses, and 86 woody species), 493 insect species (464 moths and 27 mantises), 18 bat species, 38 mammal species (20 herbivores, 6 carnivores, and 11 omnivores), and 156 bird species (35 herbivores, 105 carnivorous, 16 omnivorous). Species richness per plot varied substantially among taxonomic groups (Fig. S1). Insects exhibited the highest mean species richness (mean ± SD: 122 ± 30 species), followed by plants (64 ± 16), birds (28 ± 12), mammals (16 ± 4), and bats (14 ± 2). Species richness variability across plots was greatest for insects and birds, while bats and mammals showed narrower ranges.

GLMMs revealed significantly declining species richness of bats, birds, mammals, and insects with increasing mopane cover, whereas plant species richness showed no significant response (Table 1; Fig. 2a-e). Birds exhibited the steepest decline, with an estimated 58% reduction between the plots with the lowest and highest mopane cover, followed by mammals (30%), bats (24%), and insects (22%) (Fig. 3). Standardized effect sizes showed the same pattern of stronger negative responses in birds and mammals relative to other groups (Fig. 3). The responses of functional groups to increasing mopane cover varied markedly across taxonomic groups (Table 1; Fig. 2f-p). Species richness of plant growth forms remained largely unaffected, although grasses showed a marginally significant positive trend. In contrast, species richness of all herbivorous groups, as well as carnivorous birds and mammals and omnivorous birds, declined significantly, while carnivorous mammals and insects, and omnivorous mammals, showed no significant response.

**Figure 2.**
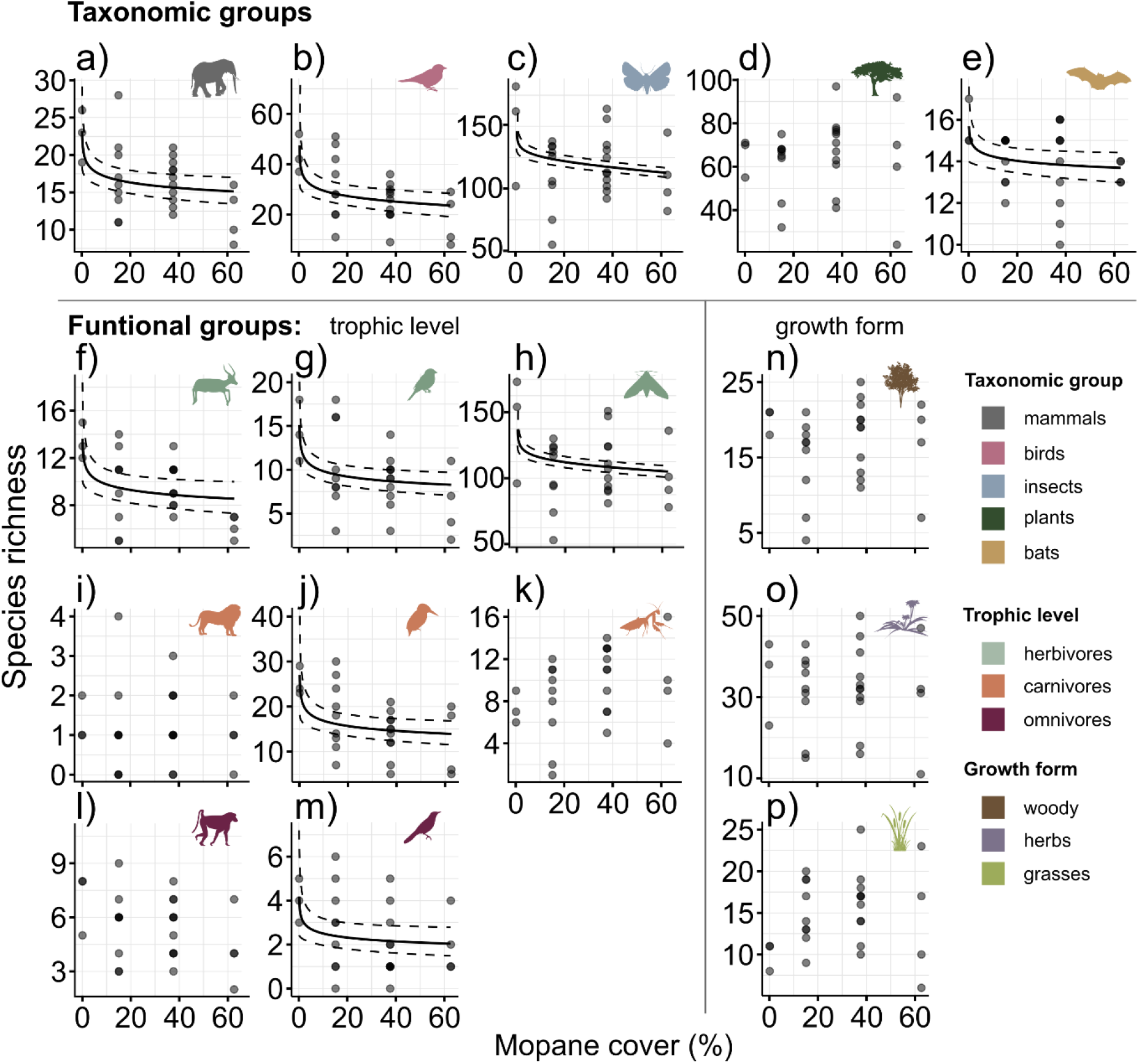
Predicted relationships between mopane cover and species richness across taxonomic and functional groups, derived from generalized linear mixed models (GLMMs). Notice, that the mopane cover values were back-transformed from the logit-transformed predictors to percentage scale for interpretability, and the visualised significant prediction lines are thus not linear. Solid lines show model predictions for groups where the effect of mopane cover was statistically significant (p < 0.05); dashed lines represent the corresponding 95% confidence intervals. The first row (a–e) shows taxonomic groups: (a) mammals, (b) birds, (c) insects, (d) plants, and (e) bats. The following rows display the corresponding functional groups of each taxon: (f–m) trophic guilds for animals (herbivores, carnivores, omnivores) and (n-p) growth forms for plants (grasses, herbs, woody species). See Table 1 for model outcomes.

**Figure 3.**
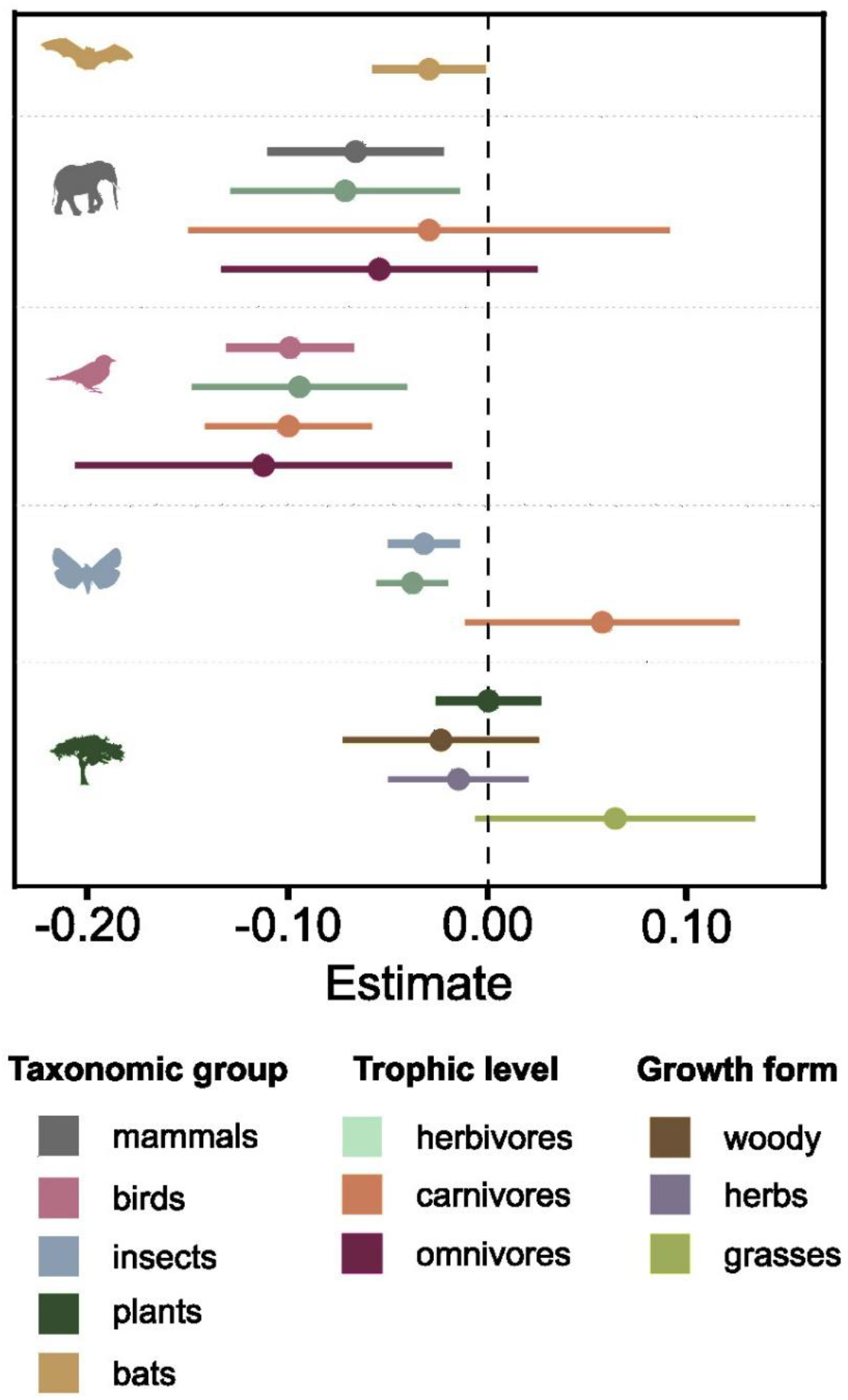
Standardized regression coefficients (β) for the effects of mopane cover on species richness across taxonomic and functional groups, estimated with generalized linear mixed models (GLMMs). Negative values indicate declines in richness with increasing mopane cover, and positive values indicate increases. Error bars show 95% confidence intervals.

Ordination analyses revealed variable effects of mopane cover on the community composition of individual taxonomic groups (Fig. 4; Table 2). Among plants, communities of herb and woody species showed marginally significant shifts. Among animals, bats, birds, and mammals, but not insects, exhibited significant changes in community composition along the mopane gradient. Among functional groups, only herbivorous mammals and omnivorous birds displayed significant community shifts in response to mopane cover.

**Table 2.**
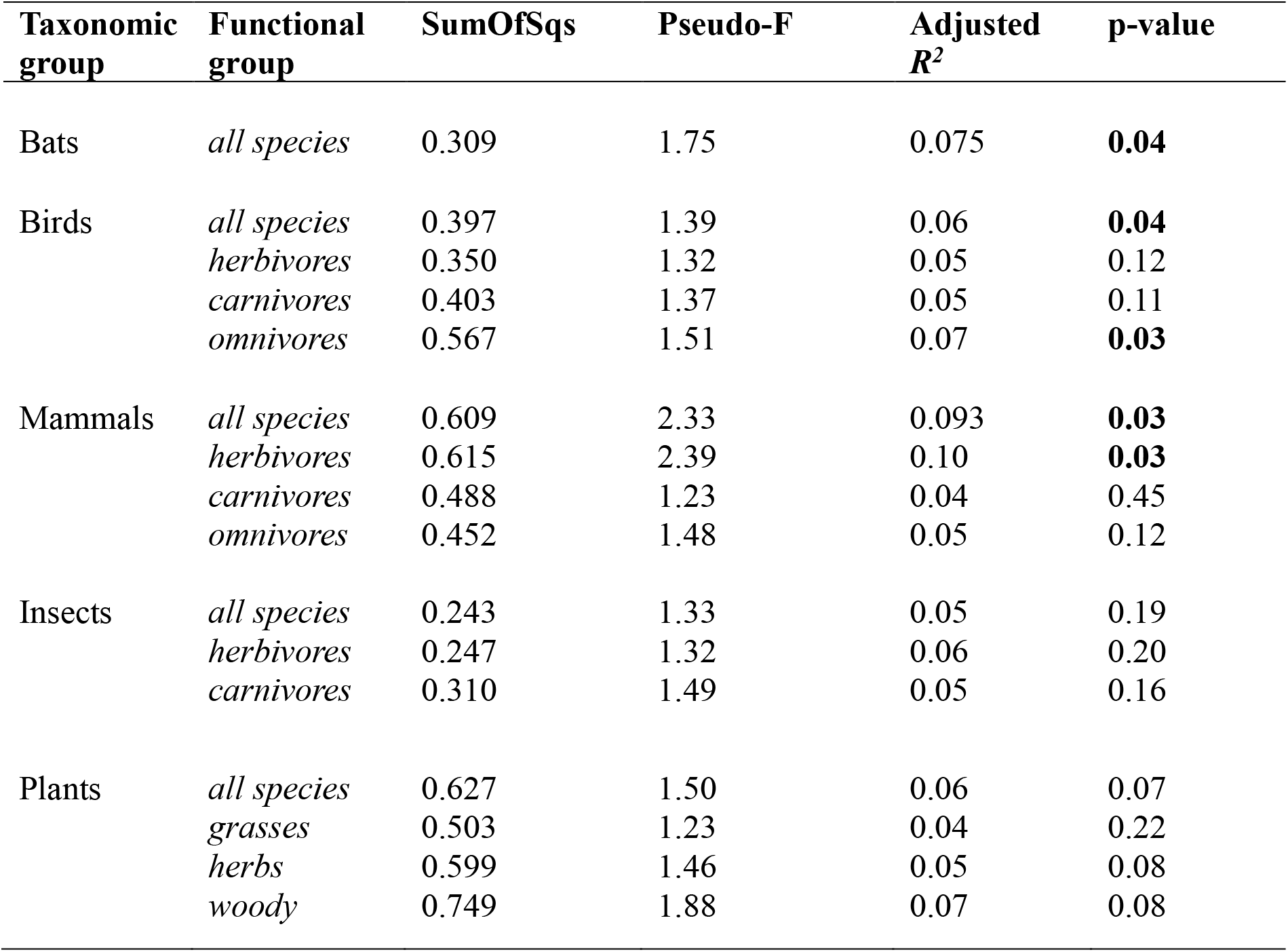
Results of distance-based redundancy analyses (db-RDA) testing the effect of mopane cover (logit-transformed) on community composition (Bray-Curtis dissimilarity) of taxonomic groups and their respective functional groups in Kruger National Park, South Africa. For each model the table reports the sum of squares attributed to mopane cover (SumOfSqs), the Pseudo-F statistic, the adjusted coefficient of determination (adjusted *R*^*2*^), and the p-value (9999 permutations, stratified by triplet). Significant compositional shifts (p < 0.05) are highlighted in bold. See Fig. 3 for significant results visualisation.

**Figure 4.**
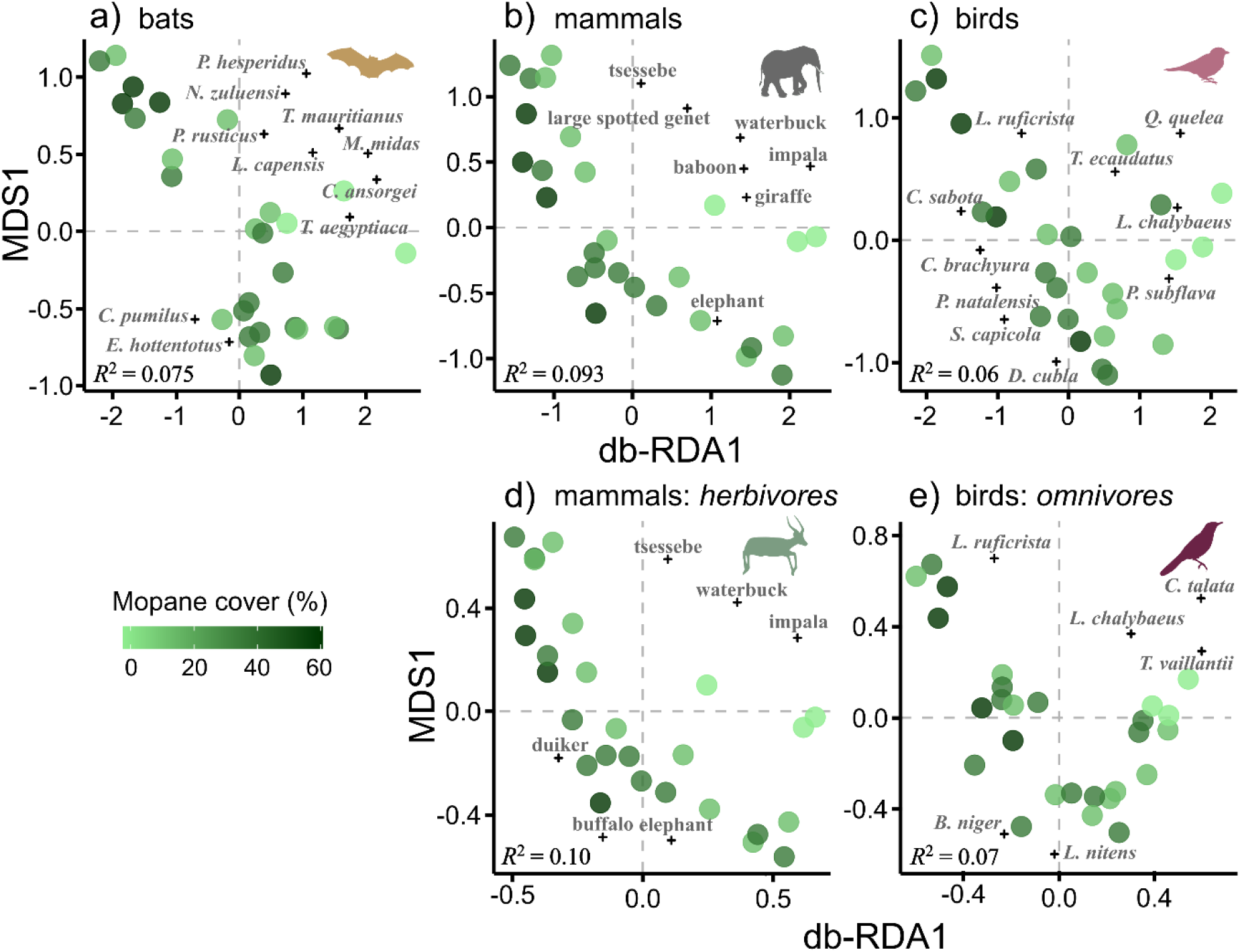
Distance-based redundancy analysis (db-RDA) ordinations illustrating significant shifts in community composition (Bray-Curtis dissimilarity) along the gradient of mopane cover in Kruger National Park, South Africa. Panels (a–c) show effects on taxonomic groups: a) bats, b) mammals, and c) birds, whilst panels (d–e) show functional groups: (d) mammal herbivores and (e) bird omnivores. Each point represents a sampling plot, colored by mopane cover percentage (darker hues indicate higher dominance). Species with strong contributions to db-RDA1 are labelled using abbreviated scientific names (a, c, e) or common names (b, d). The corresponding full species names are listed in Table S1. The proportion of community variation explained by mopane cover (adjusted *R*^*2*^) is shown in each panel. Complete results of db-RDA are provided in Table 2.

## Discussion

Our study demonstrated that increasing dominance of *Colophospermum mopane* in African savannas was generally associated with reduced biodiversity. Species richness declined in multiple vertebrate groups and insects, with the steepest reductions in birds, while overall plant richness remained unchanged and grasses showed a weak positive trend. Vertebrate assemblages (birds, mammals, bats) also exhibited modest but significant compositional shifts along the mopane gradient, whereas insect communities lost species without detectable systematic changes in composition. These patterns are consistent with previous taxon-specific observations on effects of mopane dominance (e.g. Timberlake, 1995; Khavhagali & Ligavha-Mbelengwa, 2009; Georginah & Maanda, 2015), and with effects reported for other monodominant woody species such as *Acacia mellifera* and *Terminalia sericea* (Wiegand et al., 2006; Nakafeero et al., 2007). More broadly, they align with evidence that woody plant encroachment simplifies biodiversity in tropical grassy biomes (Eldridge et al., 2011; Belay et al., 2013).

Together, these patterns indicated that mopane functions as a strong ecological filter across trophic levels. Mopane’s adaptations simplify understorey structure by reducing light and water availability, accumulating slowly decomposing litter, and promoting low multi-stem coppice (Smit & Rethman, 1998; Mlambo & Mwenje, 2010; Mlambo & Mapaure, 2006; Smallie & O’Connor, 2000; Makhado et al., 2014), thereby limiting host plants and altering food webs in Afrotropical savannas (Sankaran et al., 2005; Parr et al., 2014). This bottom-up filtering is consistent with declines in insects, whose diversity is known to be closely tied to vegetation composition and structure (Scherber et al., 2010; Parker et al., 2023; Gaona et al., 2025), and with consequent declines in insectivorous vertebrates. Birds and bats exhibited the clearest biodiversity reductions, mirroring regional evidence that woody encroachment erodes open-savanna assemblages of both groups, which depend on structurally diverse foraging habitats and abundant insect prey (Sirami et al., 2009; Sirami & Monadjem, 2012; Shapiro et al., 2020). Mammal responses were most apparent in herbivores, with limited effects on carnivores and omnivores, further confirming that bottom-up processes shaped community responses, despite mopane’s value as dry-season forage for some browsers (Ben-Shahar, 1996; Styles & Skinner, 2000; Makhado et al., 2012).

These associations between higher mopane cover and reduced richness and altered community composition across taxa support the growing consensus that Afrotropical savanna biodiversity depends on spatiotemporal heterogeneity, including open habitats, rather than maximal tree cover (Parr et al., 2014; Newbold et al., 2015; Bond et al., 2019). Such changes are of particular concern because vertebrates and insects provide key ecosystem functions such as herbivory, seed dispersal, pollination, scavenging, and herbivore suppression. Mopane dominance therefore likely affects not only species composition but also ecosystem functioning. The shift towards communities with fewer species and a higher proportion of mopane-tolerant specialists is unlikely to substitute for functionally diverse open-savanna assemblages (Parr et al., 2014) and probably reduces the resilience of savanna ecosystems to climate change and other disturbances (Oliver et al., 2015).

Therefore, our results carry important implications for conservation management. Although mopane is native to southern African savannas, its local dominance may be strengthened by general drivers of broader woody encroachment, including altered fire regimes and changes in large mammal populations (Craigue et al., 2010; Stevens et al., 2016; Venter et al., 2018). Where conservation objectives prioritise open-savanna biodiversity, targeted reductions of mopane cover may restore habitat structure and diversity (Wedel et al., 2024). In such cases, conservation outcomes will depend on maintaining landscape heterogeneity and avoiding extensive, uninterrupted mopane stands. Management interventions should therefore aim to maintain or restore landscape mosaics in which mopane-dominated stands are interspersed with mixed woodland and open savanna. Fire and herbivory management are particularly important, as frequent burning and intense elephant browsing can convert tall mopane woodlands into low coppice shrublands of limited biodiversity value (Gandiwa & Kativu, 2009; O’Connor et al., 2024), whereas prolonged fire exclusion may promote canopy closure and further reduce habitat heterogeneity. In large protected areas such as KNP, supporting functionally diverse herbivore populations and natural disturbance processes is likely the most effective way to sustain diverse vegetation structures (Smallie & O’Connor, 2000; Mlambo & Mapaure, 2006). In smaller reserves or fragmented landscapes where natural processes are suppressed, targeted measures such as selective thinning, prescribed burning, or controlled grazing may be necessary to prevent excessive mopane dominance, structural simplification, and potentially lower associated biodiversity.

Future projections of mopane expansion under climate change (Kapuka et al., 2022; Jinga et al., 2023) warrant particular caution, as they are likely to intensify the ongoing process of woody encroachment in African savannas (Venter et al., 2018; Bond et al., 2019). Mopane’s considerable value as fuel, timber, fodder, and for mopane-worm harvest provides strong incentives for its active support by local communities. It is even promoted in savanna restoration projects, particularly outside protected areas, which together may further amplify its advantage under climate change (Jinga et al., 2023). However, deliberate planting or promotion of mopane for its economic value, should be carefully reconsidered. Coppiced mopane shrublands are difficult to reverse once established (Timberlake, 1995; Teketay et al., 2018), and expanding mopane dominance risks accelerating woody encroachment, homogenising open-biome habitats, and further reducing biodiversity and ecosystem services characteristic of savanna ecosystems (Parr et al., 2014; Nerlekar & Veldman, 2020; Bond et al., 2019). Conservation strategies should therefore prioritise maintaining mosaics of open and mixed woodland savanna, limiting mopane expansion outside its current distribution, protecting open habitat refugia, and critically evaluating mopane-based livelihood or planting initiatives against their biodiversity consequences and long-term reversibility.

## Acknowledgements

We are grateful to Pavla Šonská for help in the field; to Desmond Mabaso, Herman Ntimane, and Annoit Mashele for accompanying us in the field and keeping us safe; to Samantha Mabuza, Sharon Thompson, and Patricia Khoza for their assistance with arranging permits and other logistics in KNP; and to Freerk Molleman for comments on an earlier manuscript draft. We thank SANParks for their support (project PYSK 1432), noting that the results do not reflect the opinions of SANParks but the authors alone. We used ChatGPT 4o and 5 (OpenAI) language model for English proofreading. This study was funded by the Czech Science Foundation (18-18495S and 21-24186M). P.Py., J.Č., M.H. and K.P. were also supported by the long-term research development project RVO 67985939 (Czech Academy of Sciences).

## Author Contribution

FPG, PPy, DS, and RT conceived the study idea; PPy, DS, LCF, MH, SM, KP, and RT designed the study and established the study plots within the MOSAIK project; all authors sampled, recorded, and/or identified focal organisms; FPG, MH, and RT performed analyses; FPG and RT interpreted results, prepared visualisations, and wrote the first draft; all authors contributed to writing and approved the final manuscript.

## Notes

### Competing Interest Statement

The authors have declared no competing interest.

