## Supplementary Material for "Effects of a native, dominant tree, *Colophospermum mopane*, on diversity of plants, insects, and vertebrates in South African savannas"


Fernando P. Gaona[1,2,*], Tomáš Albrecht[3,4], Jan Čuda[5], Sylvain Delabye[1,2], Llewellyn C. Foxcroft[6,7], Valeriy Govorov[3], Martin Hejda[5], Ivan Horáček[3], Sandra MacFadyen[8,9], Pavel Potocký[2], Klára Pyšková[5,1], Vladimír Remeš[1,10], Ondřej Sedláček[1], Markéta Staňková[3], David Storch[11,1], Petr Pyšek[5,1], Robert Tropek[1,2,*]

[1]*Department of Ecology, Faculty of Science, Charles University, Prague, Czechia*
[2]*Institute of Entomology, Biology Centre, Czech Academy of Sciences, Ceske Budejovice, Czechia*
[3]*Department of Zoology, Faculty of Science, Charles University, Prague, Czechia*
[4]*Institute of Vertebrate Biology, Czech Academy of Sciences, Brno, Czechia*
[5]*Czech Academy of Sciences, Institute of Botany, Department of Invasion Ecology, Průhonice, Czechia*
[6]*Scientific Services, South African National Parks, Skukuza, South Africa*
[7]*Centre for Invasion Biology, Department of Botany and Zoology, Stellenbosch University, Matieland, South Africa*
[8]*Mathematical Biosciences, Stellenbosch University, Department of Mathematical Sciences, Stellenbosch, South Africa*
[9]*National Institute for Theoretical and Computational Sciences, Stellenbosch University, Matieland, South Africa*
[10]*Department of Zoology, Faculty of Science, Palacky University, Olomouc, Czechia*
[11]*Center for Theoretical Study, Charles University, Prague, Czechia*

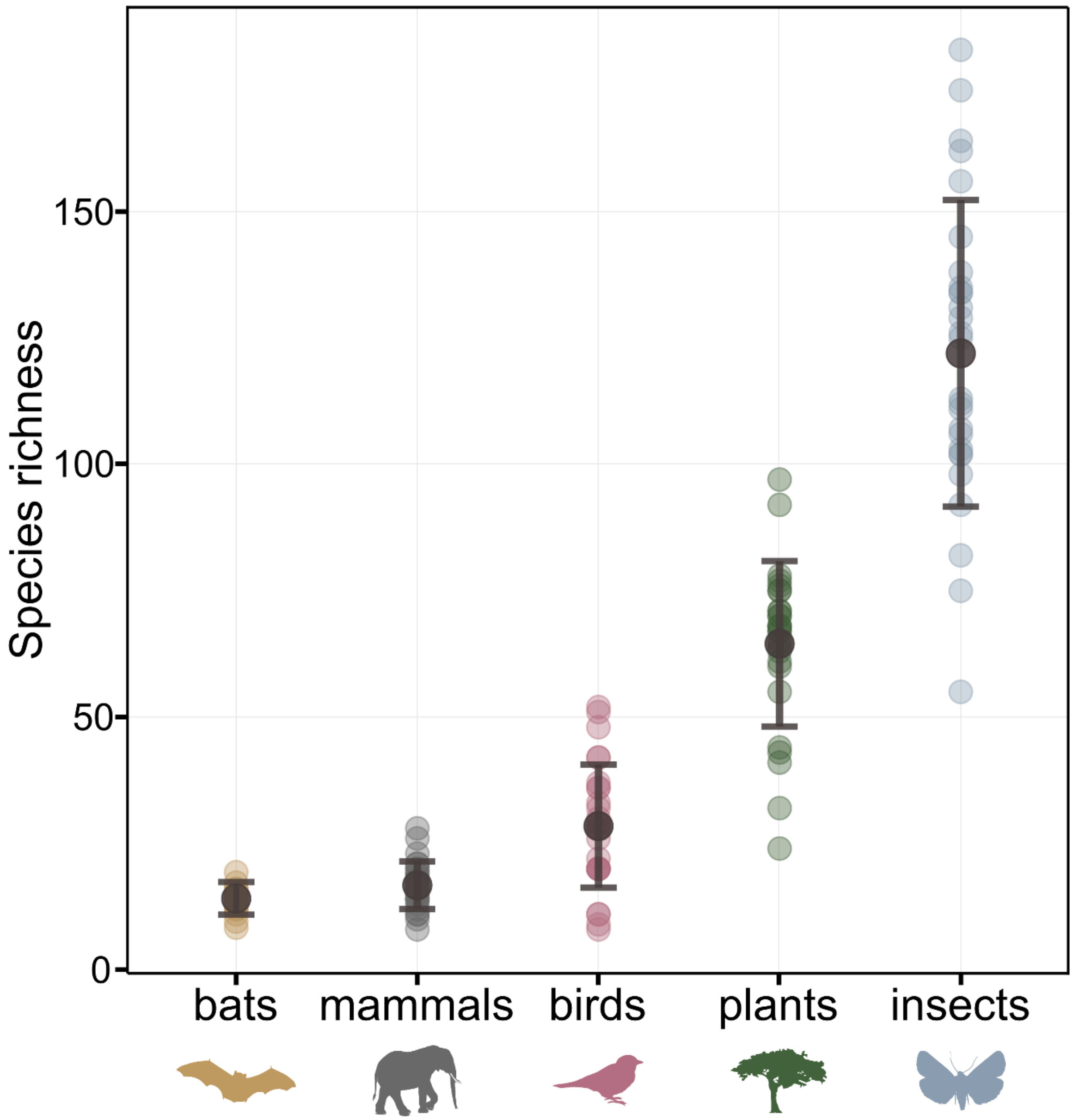


**Figure S1.** Mean species richness across six taxonomic groups: bats, mammals, birds, plants and insecs. Each point represents the observerd species richness in one of the 27 sampled plots within the study area. Larger dark circles indicate the mean species richness for each group, and vertical error bars represent the standard deviation (SD) across plots.

**Table S1.** Label used in Figure 4 (either common name or abbreviated scientific name), and full scientific name of each species contributing most strongly to the distance-based redundancy analysis (db-RDA) ordinations of community composition.

| Taxonomic group | Abbreviation/ Common name | Scientific name |
|---|---|---|
| bats | *C. ansorgei* | *Chaerephon ansorgei* |
| | *C. pumilus* | *Chaerephon pumilus* |
| | *E. hottentotus* | *Eptesicus hottentotus* |
| | *L. capensis* | *Laephoptis capensi* |
| | *M. midas* | *Mops midas* |
| | *N. zuluensi* | *Neoromicia zuluensi* |
| | *P. herperidus* | *Pipistrellus hesperidus* |
| | *P. rusticus* | *Pipistrellus rusticus* |
| | *T. aegyptiaca* | *Tadarida aegyptiaca* |
| | *T. mauritianus* | *Taphozous mauritianus* |
| birds | *B. niger* | *Bubalonis niger* |
| | *C. bracyura* | *Cameroptera bracyura* |
| | *C. sabota* | *Calendulauda sabota* |
| | *C. talata* | *Cinnyris talata* |
| | *D. cubla* | *Dryopcopus cubla* |
| | *L. chalybaeus* | *Lamprotornis chalybaeus* |
| | *L. nitens* | *Lamprotornis nitens* |
| | *L. rufricrista* | *Lophotis rufricrista* |
| | *P. natalensis* | *Pternistis natalensis* |
| | *P. subflava* | *Prinia subflava* |
| | *Q. quelea* | *Quelea quelea* |
| | *S. capicola* | *Streotopelia capicola* |
| | *T. ecaudatus* | *Terathopius ecaudatus* |
| | *T. vaillantii* | *Trachyphonus vaillantii* |
| mammals | baboon | *Papio ursinus* |
| | elephant | *Loxodonta africana* |
| | duiker | *Sylvicapra grimmia* |
| | giraffe | *Giraffa giraffa* |
| | impala | *Aepyceros melampus* |
| | large spotted genet | *Genetta tigrina* |
| | tsessebe | *Damaliscus lunatus* |
| | waterbuck | *Kobus ellipsiprymnus* |
